# AFEchidna is an R package for genetic evaluation of plant and animal breeding datasets

**DOI:** 10.1101/2021.06.24.449740

**Authors:** Weihua Zhang, Ruiyan Wei, Yan Liu, Yuanzhen Lin

## Abstract

Progeny tests play important roles in plant and animal breeding programs, and mixed linear models are usually performed to estimate variance components of random effects, estimate the fixed effects (Best Linear Unbiased Estimates, BLUEs) and predict the random effects (Best Linear Unbiased Predictions, BLUPs) via restricted maximum likehood (REML) methods in progeny test datasets. The current pioneer software for genetic assessment is ASReml, but it is commercial and expensive. Although there is free software such as Echidna or the R package sommer, the Echidna syntax is complex and the R package functionality is limited. Therefore, this study aims to develop an R package named AFEchidna based on Echidna software. The mixed linear models are conveniently implemented for users through the AFEchidna package to solve variance components, genetic parameters and the BLUP values of random effects, and the batch analysis of multiple traits, multiple variance structures and multiple genetic parameters can be also performed, as well as comparison between different models and genomic BLUP analysis. The AFEchidna package is free, please email us to get a copy if reader is interested for it. The AFEchidna package is developed to expand free genetic assessment software with the expectation that its efficiency could be close to the commercial software.

## Introduction

Mixed linear models are widely used in the analysis in the progeny test data of plants and animals [1–4]. Mixed linear models are linear models with a combination of fixed and random effects to explain the degree of variation in interest traits, such as milk yield in cows or volume growth in forest trees. Breeders are often interested in predicting the future performance of a particular genotype of animal or crop in an environment, and treat the underlying genetic factors affecting the target trait as random effects. By contrast, breeders are less interested in site-specific effects or experimental replication within the site, which are generally treated as fixed effects [5]. Therefore, mixed linear models are well suited for genetic data analysis. Nowadays, most software uses the Restricted Maximum Likelihood (REML) method to estimate the variance components of random effects, and then estimate the fixed effects and predict the random effects.

Progeny tests are widely designed in almost all genetic improvement projects in plants and animals. In the analysis of progeny test data, breeders typically focus on the additive effects. In a given population, an additive effect could be stably inherited from the parents to the offspring, so the ratio of additive genetic variance to phenotypic variance is defined as the narrow heritability (h2) of a target trait. The amount of additive variance and heritability have a great influence on the genetic gain [6,7]. Current progeny test datasets usually have pedigree data, phenotypic data and marker data. Professional genetic analysis software can form A matrix from pedigree data or G matrix from marker data using in the variance component estimation and BLUP of random effects [8,9]. Such genetic analysis software includes ASReml [10], Echidna [11], SAS[12], BLUPF90 [13], and R packages sommer [14], breedR [15], etc. Table 1 lists the advantages and disadvantages of common genetic analysis software. For forest experiments, due to the complexity of test environmental factors, software is needed to fit the complex variance structure [16–18]. At present, the commercial software ASReml is recognized as the pioneer software for plant genetic assessment, but it is expensive. Although the breedR package is developed for forest experiment, its version is old and the variance structure type is few. Echidna is a free software developed in 2018 by Professor Gilmour, the main developer of ASReml [11]. It also uses REML method to estimate parameter values, and its syntax and function is very close to that of ASReml. It is the most powerful free software for animal and plant genetic assessment, but its usage is a little complicated, which may be difficult for ordinary users. Therefore, the purpose of this paper is to provide AFEchidna, an R package based on Echidna software, and demonstrate how to use a mixed linear model to generate solutions for variance components, genetic parameters, and random effects BLUPs using a half-sib dataset. In addition, the AFEchidna package also provides batch analysis of different traits and different variance structures, batch calculation of different genetic parameters, as well as calculation of genomic relationship matrix and genomic BLUP analysis.

**Table 1.**
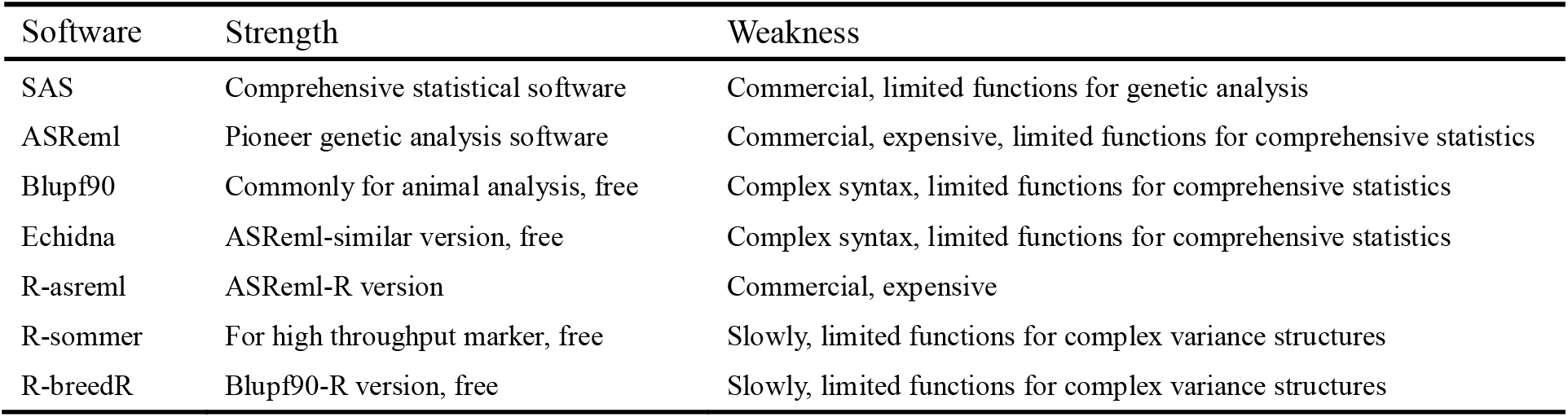
Comparison of the commonly used genetic assessment software

## Materials and methods

### Development strategy of AFEchidna package

The stand-alone version of Echidna performs mixed linear model analysis using the .es file and generates a series of result files. Due to the complicated syntax and the result file sets, also the limitation of data processing and visualization, Echidna maybe not suitable for the general users. Therefore, we use R language 4.0.2 to build the R package AFEchidna based on Echidna stand-alone version (V1.52). Through this package, users can not only solve the variance component, genetic parameters and random effect BLUP values, but also carry out the batch analysis of multiple traits, multiple variance structures and multiple genetic parameters, as well as comparisons between different models and genomic BLUP analysis.

Figure 1 shows the workflow of the AFEchidna package. The AFEchidna package combines the primary function echidna() with the command file .es0, phenotypic data, pedigree and even marker relationship matrix files for mixed linear model analysis, and saves the running results as a R object. Then, we use function Var() to extract, and coef() to obtain the model solution, as well as predict() for the model predicted values and pin() for the genetic parameter values. We also design update() for running the new model and model.comp() to compare different models. Table 2 lists the summary of the dependent R packages, descriptions, and simple usage for the listed functions in AFEchidna.

**Figure 1.**
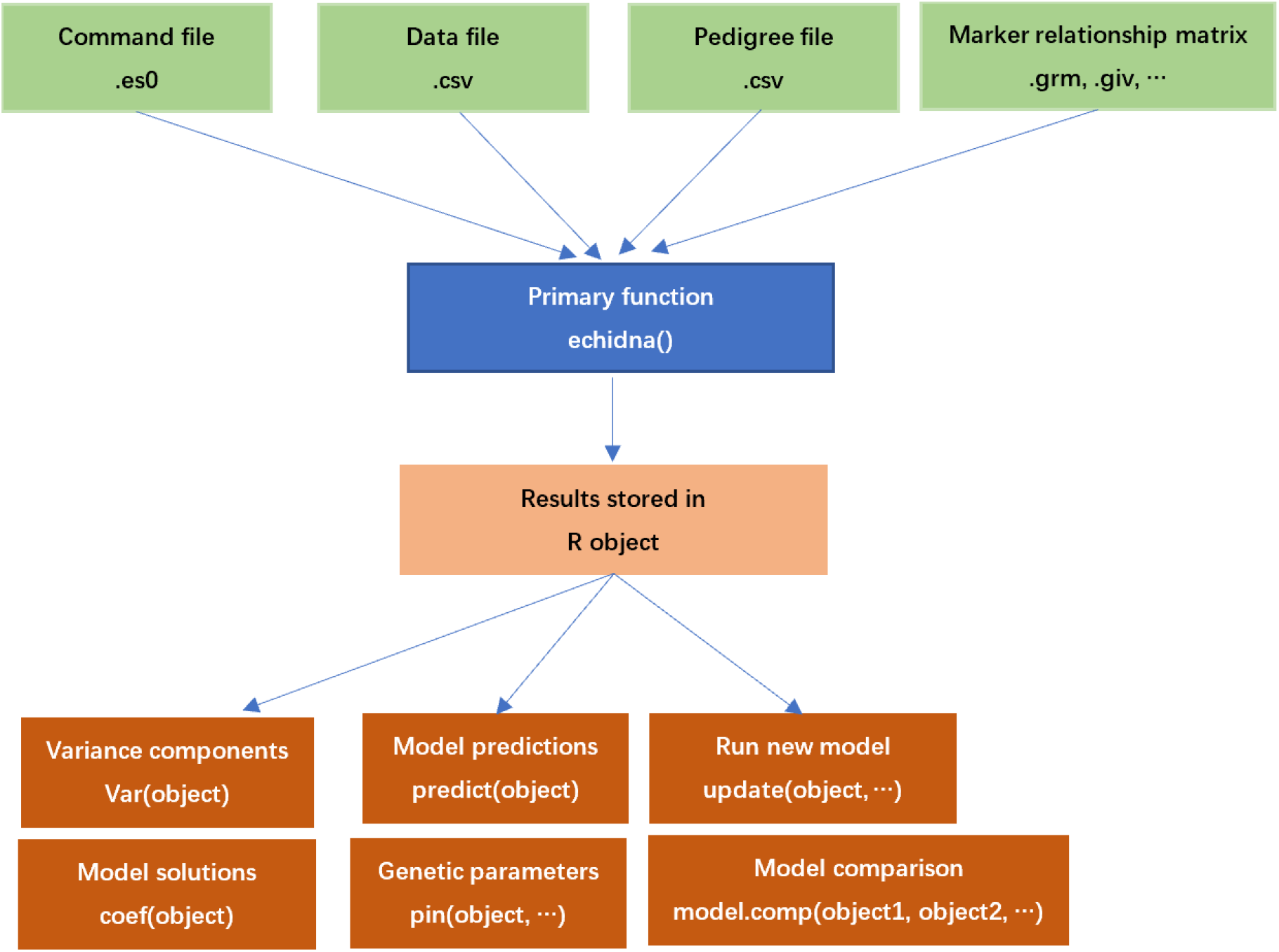
The workflow of AFEchidna package

**Table 2.** The summary of listed functions in AFEchidna package

| Function | Dependent package | Description | Simple usage |
| --- | --- | --- | --- |
| checkPack | utils | Check dependent packages | checkPack() |
| coef | dplyr | Model equation solutions | coef(object) |
| echidna | dplyr, tidyr, readr, stringr, reshape | Specify mixed linear model | echidna(fixed, random, residual, es0.file, ...) |
| GenomicRel | GeneticsPed | Generate genomic relationship matrix | GenomicRel(marker,option,ped,...) |
| get.es0.file |  | Generate es0 file | get.es0.file(dat.file, es.file, ...) |
| model.comp | stats | Compare different models | model.comp(object1, object2,LRT) |
| pin | dplyr, msm | Calculate genetic parameters | pin(object, ...) |
| predict | readr | Model predictions | predict(object) |
| plot | dplyr, ggplot2 | Model diagnose plots | plot(object) |
| update |  | Run new model | update(object, ...) |

Install the AFEchidna package from the local file AFEchidna_1.0.2.zip, and run AFEchidna:: CheckPack ()in R language to check whether the R dependent package is installed or not. If not, it will be installed automatically. The AFEchidna package is free to the public, please email to to get a copy. The AFEchidna package is for academic research only and not for commercial use.

### Example case

The dataset is from Example 4.5 in Reference [4]. This dataset has 36 half-sib families of a pine tree from four provenances. The experimental design was a randomized complete block design with five replications and from two to six trees measured within each plot.

Taking tree height as the target trait, the mixed linear model is assumed as follows:

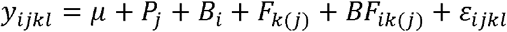

Where, *cy_ijkl_* is the *l*th observation of the *i*th block, *j*th provenance and *k*th family; *μ* is the overall mean; *P_j_* is the fixed provenance effect; *B_i_* is the random *i*th block effect, 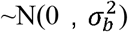; *F_k(j)_* is the random *k*th female effect within its provenance group, 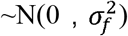;, *BF_ik(j)_* is the random block by female interactions, 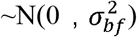; *ε_ijkl_* is the random residual effect, 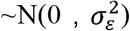.

Individual-tree narrow-sense heritabilities were estimated as follows:

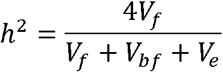

where, *V_f_* is the female variance, *V_bf_* is the block by female interaction variance, *V_e_* is the residual variance.

The breeding value and its accuracy of female were estimated as follows:

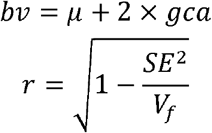

Where, *bv* is female breeding value, *gca* is female general combining ability, *μ* is the overall mean, *r*, is the accuracy of female breeding value, *SE* is the standard error of female general combined ability, *V_f_* is the female variance.

The above statistical analysis was implemented by the AFEchidna and compared with the results of ASReml software [4].

## Results

### Variance component and heritability estimates

The variance components estimated by the AFEchidna package and the ASReml software showed that the estimated values of the four variance components including the maternal variance Vf and their standard errors were all consistent (Table 3). The results of genetic parameters, such as heritability, were also completely consistent between AFEchidna and the ASReml (Table 4). These results suggest that the AFEchidna package is equivalent to the ASReml in estimating the variance components of mixed linear models.

**Table 3.** The estimation of variance components from AFEchidna and ASReml

| Software | Variance components |  |  |  |  |  |  |  |
| --- | --- | --- | --- | --- | --- | --- | --- | --- |
| | $V_b$ | $V_b.se$ | $V_f$ | $V_f.se$ | $V_{bf}$ | $V_{bf.se}$ | $V_e$ | $V_e.se$ |
| AFEchidna | 0.107 | 0.090 | 0.190 | 0.084 | 0.198 | 0.086 | 2.527 | 0.132 |
| ASReml | 0.107 | 0.090 | 0.190 | 0.084 | 0.198 | 0.086 | 2.527 | 0.132 |

Note: $V_b$ , $V_f$ , $V_{bf}$ , $V_e$ is the variance components of block, female, block by female and residual error , respectively; se is the standard error.

**Table 4.** The estimation of genetic parameters from AFEchidna and ASReml

| Software | Genetic parameters |  |  |  |  |  |
| --- | --- | --- | --- | --- | --- | --- |
| | $V_a$ | $V_a.se$ | $V_p$ | $V_p.se$ | $h^2$ | $h^2.se$ |
| AFEchidna | 0.758 | 0.336 | 2.914 | 0.150 | 0.260 | 0.110 |
| ASReml | 0.758 | 0.336 | 2.914 | 0.150 | 0.260 | 0.110 |

Note: $V_a$ , $V_p$ , $h^2$ is the additive variance, phenotypic variance and heritability , respectively; se is the standard error.

### Solutions of fixed effects

The fixed effect includes population mean (μ) and provenance (Prov), and their unbiased estimates are shown in Table 5. In AFEchidna, the fixed population mean is 0, and the effect values at each level of provenance are directly given. In ASReml, the effect value of the first level of the provenance was fixed as 0, and then the population mean and other level values of the provenance were calculated. Thus, AFEchidna and ASReml have slightly different methods for solving fixed effects and slightly different results. However, the provenance effect values obtained by ASReml are relative values, and when they are added to the population mean, they are basically consistent with the results of AFEchidna.

**Table 5.** The solutions of fixed effects

| software | Fixed effect | Level | Solution | Standard error |
| --- | --- | --- | --- | --- |
| AFEchidna | $\mu$ | 1 | 0.000 | 0.000 |
|  | Prov | 10 | 11.512 | 0.362 |
|  | Prov | 11 | 9.846 | 0.228 |
|  | Prov | 12 | 10.888 | 0.221 |
|  | Prov | 13 | 10.292 | 0.233 |
| ASReml | $\mu$ | 1 | 11.510 | 0.362 |
|  | Prov | 10 | 0.000 | 0.000 |
|  | Prov | 11 | -1.666 | 0.374 |
|  | Prov | 12 | -0.624 | 0.370 |
|  | Prov | 13 | -1.220 | 0.377 |

Note: $\mu$ , Prov is overall mean and provenance, respectively.

### The breeding value and its accuracy of female

As shown in Table 6, AFEchidna and ASReml obtained the same results of the general combining ability and standard error of females. Due to their slightly different estimated population means, the estimated breeding values of females were also slightly different, but the correlation between them reached 0.999(*P*<.001, Figure 2). The accuracy of the breeding values was also consistent.

**Figure 2.**
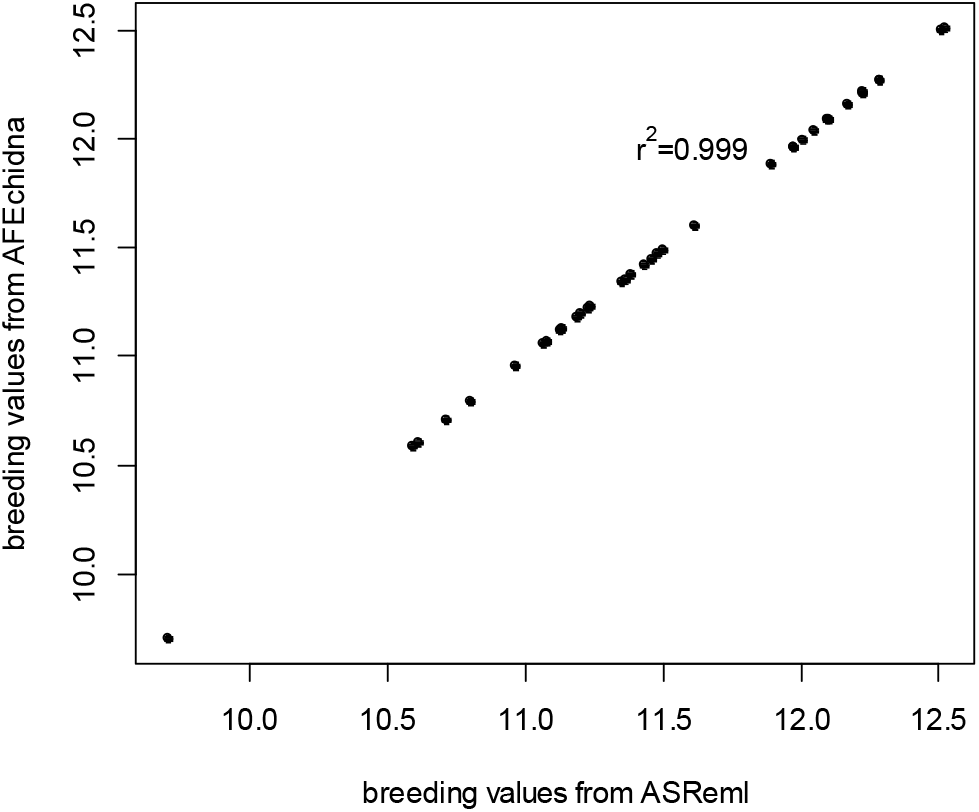
Scatter plot of female breeding values from ASReml and AFEchidna

**Table 6.**
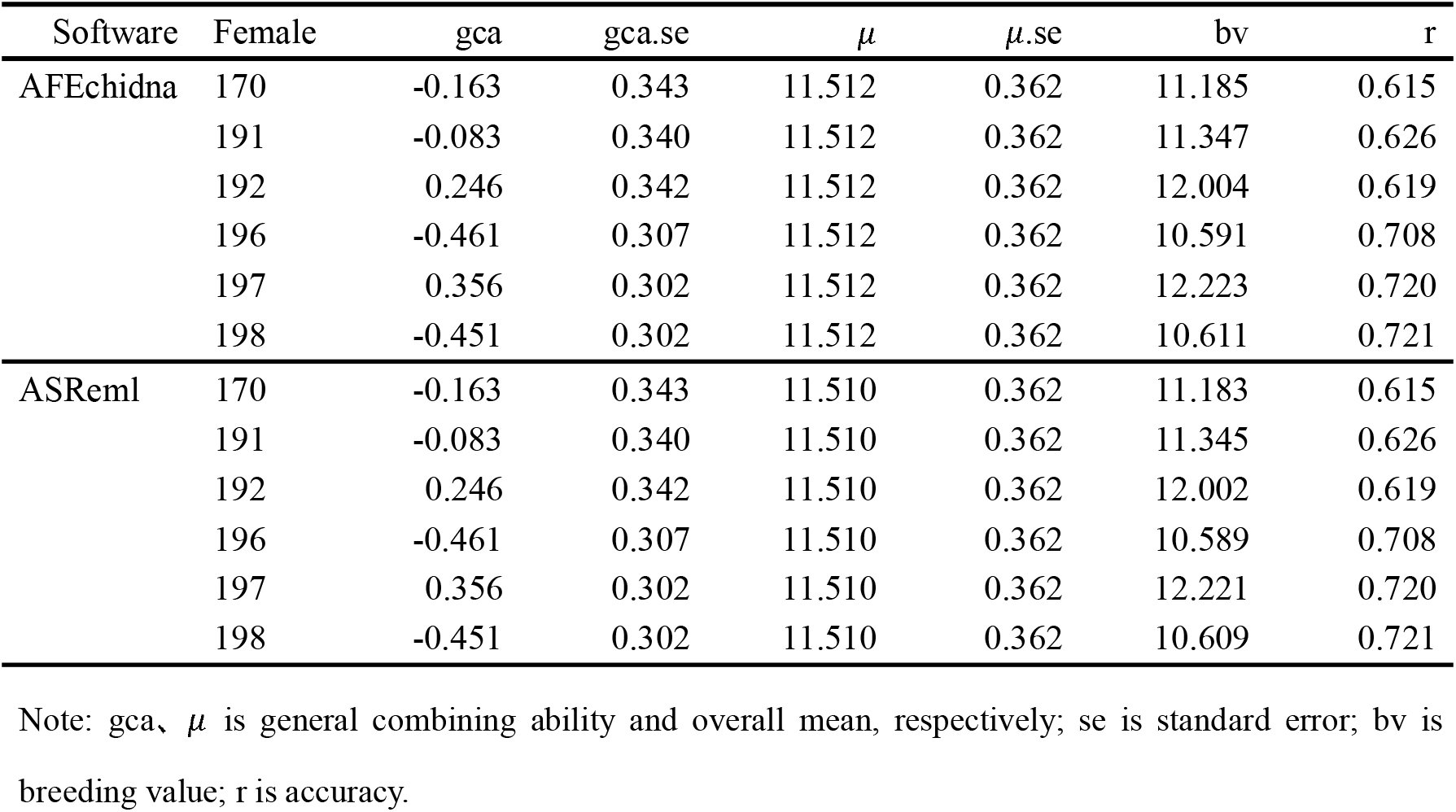
The first six female breeding values and their accuracy

### Significant test for random effects

The mixed linear model in this study was regarded as a full model, and its random effects included block effect, female effect and block×female effect. One of them was successively deleted as a reduced order model, and then LRT method was used to test the significance of the corresponding random effects [4]. The model.comp() function in the AFEchidna package can implement LRT test. The LRT test results (Table 7) revealed that block effect, female effect and block×female effect all had extremely significant effects on tree height (*P*<0.01).

**Table 7.** Significant test for random effects

| Model | LikelihoodLog | LRT value | *P* value | Significant level |
| --- | --- | --- | --- | --- |
| Full | -934.120 |  |  |  |
| No Block | -940.640 | 13.040 | 0.0002 | *** |
| No Female | -939.820 | 11.400 | 0.0004 | *** |
| No Block×Female | -937.890 | 7.540 | 0.0030 | ** |

Note:\*\*\* and\*\* indicate $P < 0.001$ and $P < 0.01$ , respectively.

### batch analysis of single trait

The primary function echidna() in the AFEchidna package can also perform batch analysis for multiple traits, through the batch=TRUE parameter. Batch analysis can be used not only for a single trait, but also for multiple traits. In this study, the same mixed linear model was used to conduct batch analysis of single trait for tree height, diameter and volume. The variance component results were shown in Table 8.

**Table 8.** The estimation of variance components from single trait batch analysis

| Trait | Variance components |  |  |  |  |  |  |  |
| --- | --- | --- | --- | --- | --- | --- | --- | --- |
| | $V_b$ | $V_{b.se}$ | $V_f$ | $V_{f.se}$ | $V_{bf}$ | $V_{bf.se}$ | $V_e$ | $V_{e.se}$ |
| height | 0.107 | 0.090 | 0.190 | 0.084 | 0.198 | 0.086 | 2.527 | 0.132 |
| diameter | 0.167 | 0.198 | 0.807 | 0.415 | 0.691 | 0.501 | 16.881 | 0.877 |
| volume | 83.516 | 76.846 | 219.220 | 102.160 | 164.05 | 111.470 | 3731.500 | 193.670 |

Note: all volume values are multiplied by 1000 times.

## Discussions

We developed a R package named AFEchidna based on Echidna stand-alone version (V1.52) using R language 4.0.2. Taking a half-sib dataset as an example, the mixed linear model was used to estimate the variance component of random effects and heritability, solve the fixed effects, and estimate the breeding value and its accuracy. These results were highly consistent with those of commercial software ASReml. In addition, the significance test of random effects and the batch analysis of single trait were also demonstrated.

Wang and Tong [19] developed the R package Halfsibms, which was used to analyze the half-sib progeny data at multiple sites, and they input the data file directly for genetic analysis without model term and variance specification. Although this operation may seem convenient, it has very narrow applications. Due to the wide types of forest tree genetic tests, there are not only various types of field experiment designs, but also complex genetic material sources. A simple fixed mixed model is simply not enough to include different statistical methods, such as spatial analysis model [16], factor analysis model [17] and genome selection model [20]. Therefore, the mainstream genetic analysis software (such as ASReml, Echidna, SAS, breedR, etc.) does not specify a fixed mixed model. Based on this idea, when we developed the AFEchidna package, the flexibility that the mixed linear model could specify any testing factor and its variance structure was retained, using the parameters fixed, random and residual within the primary function echidna(), which remained the powerful function of the mixed model analysis of Echidna.

Isik et al. [4] pointed out that the commercial software ASReml used average information (AI) and sparse matrix algorithms to solve a large number of mixed model equations, faster and more efficient than the SAS Proc Mixed (Newton-Raphson algorithm). ASReml can easily handle different mating designs, different field designs, multivariate models and other analyses, due to its flexible ability for diverse variance structures. The weakness of ASReml is lack of data management, comprehensive statistics, and graphical visualization capabilities. As the free sister version of ASReml, Echidna also inherits the advantages and disadvantages of ASReml. Therefore, we developed the AFEchidna package with R language, incorporating the advantages of R language in data management, comprehensive statistics and graphical visualization. For example, the LRT test method is used to test the significance of model random terms, or to compare different models. According to the LRT rule, we write the function model.comp(). For another example, with the advancement of breeding process and the accumulation of the number of target traits, the demand for batch analysis of traits is inevitable. Although the software Echidna can handle the batch analysis of single trait (the trouble is that there are too many result files and it is difficult to extract the analysis results in batches), it cannot handle the batch analysis of multi-trait models. Thus, we run batch analysis of single trait or multiple traits using the batch parameter in the primary function echidna(), and can output the results directly. In addition, there are extra parameters batch.G and batch.R in the function echidna(), which can be used for batch analysis of various G structures and various R structures. This method is proposed by us for the first time, and has not been involved in other genetic assessment software.

In summary, the AFEchidna package not only inherits the advantages of the Echidna software, which can fit different mating design models (testing cross, nest mating, diallel mating, etc.), spatial analysis model, multivariate model, multi-site model and genomic BLUP model, but also retains the advantages of R language. The function can be programmed to batch analyze traits, test the significance of random items, etc., and can use R language for data management, comprehensive statistics and graphic visualization functions. In the future, we will introduce how the AFEchidna package fits the spatial analysis model, multivariate model, multi-site model and genomic BLUP model.

## Supplemental codes

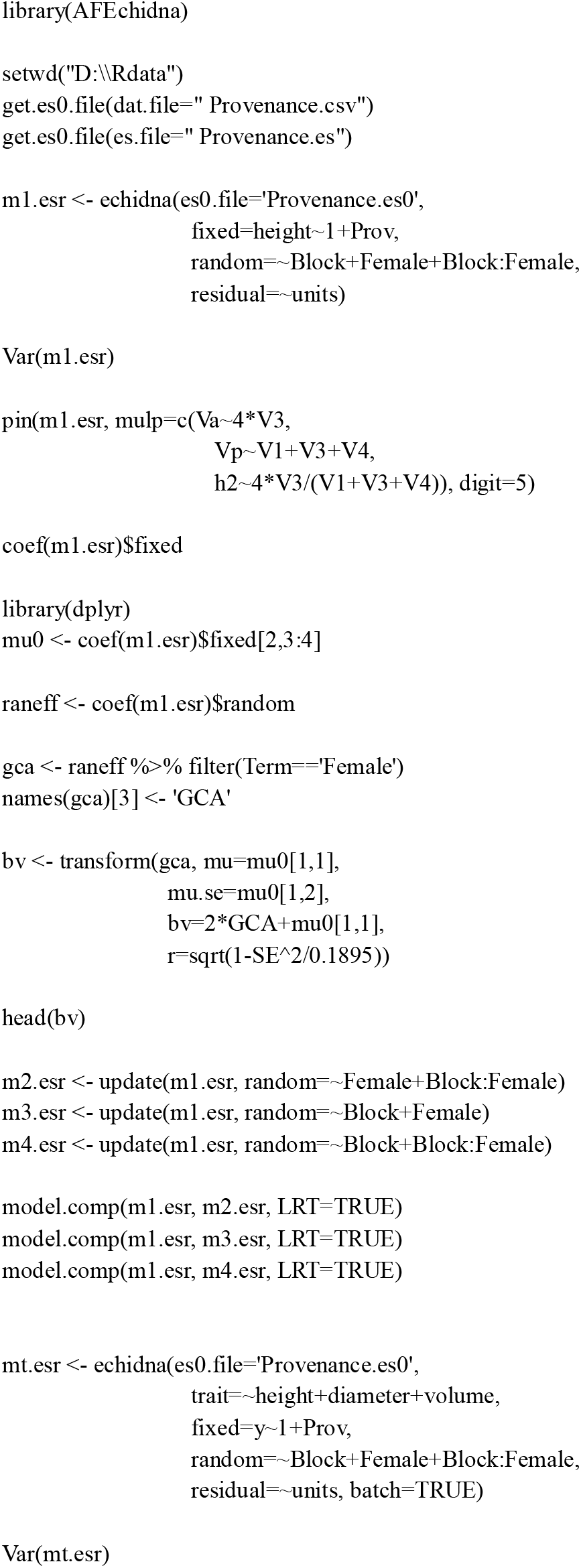

